# TIR-1/SARM1 Inhibits Axon Regeneration

**DOI:** 10.1101/2020.06.23.165852

**Authors:** Victoria Julian, Alexandra B. Byrne

## Abstract

An injured axon has two choices, regenerate or degenerate. In many neurons, the result is catastrophic axon degeneration and a failure to regenerate. To coerce the injured nervous system to regenerate, the molecular mechanisms that regulate both axon regeneration and degeneration need to be defined. We found that TIR-1/SARM1, a key regulator of axon degeneration, inhibits regeneration of injured motor axons. Loss of *tir-1* function both reduces the frequency with which severed axon fragments degenerate and increases the frequency of axon regeneration. The increased regeneration in *tir-1* mutants is not a secondary consequence of its effects on degeneration. Rather, TIR-1 carries out each of these opposing functions cell autonomously by regulating independent downstream genetic pathways. While promoting axon degeneration with the DLK-1 mitogen activated protein kinase (MAPK) signaling cascade, TIR-1 inhibits axon regeneration by activating the NSY-1/ASK1 MAPK signaling cascade. Our finding that TIR-1 regulates both axon regeneration and degeneration provides critical insight into how axons coordinately regulate the two key responses to injury, consequently informing approaches to manipulate the balance between these responses towards repair.

## Introduction

In injured neurons that are capable of repair, axons are broken into fragments with different fates. The axon segment that remains attached to the cell body regenerates and severed axon segments degenerate. However, this balance is disrupted in the central nervous system and in aged animals where axon regeneration is inhibited, resulting in a permanent loss of neuronal function (*1, 2*). Understanding the molecular mechanisms that coordinately regulate both axon regeneration and degeneration is critical to developing therapeutic approaches to repair the injured nervous system.

On the proximal side of the injury, functional axon regeneration requires the synchronized execution of multiple processes, including growth cone formation, axon guidance and synapse formation. Despite being recapitulations of well-characterized developmental processes, the precise molecular mechanisms that regulate axon regeneration remain incompletely understood. Those that have been identified are subdivided into two broad categories: intrinsic and extrinsic. Intrinsic regulators of transcription, transport, translation, and signaling are complemented and opposed by extrinsic regulators of myelin dependent inhibition and scar formation (*3–8*). Manipulating either its intrinsic or extrinsic environment is sufficient to modestly increase an injured axon’s ability to regenerate (*7, 9–11*).

On the distal side of the injury, the most notable endogenous regulator of axon degeneration is Sterile Alpha and Toll/interleukin-1 resistance (TIR) Motif Containing 1 (SARM1/dSarm), which is essential for injury-induced axon degeneration of *Drosophila* and mouse axons (*12, 13*). SARM1/dSarm promotes axon degeneration cell-autonomously, by depleting NAD^+^ and ATP, interacting with the BTB/BACK domain protein Axundead, and by interacting with mitogen-activated protein kinase (MAPK) signaling cascades, which in turn regulate calpain activation, SCG10/Stathmin 2 proteolysis, mitochondrial dysfunction, cytoskeletal disruption, and axon fragmentation (*14–21*). Whether the *C. elegans* SARM1/dSarm homolog, TIR-1, also regulates injury-induced axon degeneration is not known; however, in the absence of injury, SARM1, dSarm, and TIR-1 promote disease- and age-associated neurodegeneration (*22–25*). Despite the prominent role of SARM1/dSarm in the injury response, its function is currently understood to be restricted to the severed axon fragment distal to the injury site.

Here we report that not only does TIR-1/SARM1 promote injury-induced axon degeneration, it also inhibits axon regeneration. Specifically, in response to injury, TIR-1 functions with the NSY-1/ASK1 MAPK signaling cascade and the terminal transcription factor DAF-19/RFX to inhibit regeneration of the proximal axon fragment. In parallel, TIR-1 functions independently from NSY-1/ASK1 signaling and with the DLK-1/DLK MAPK pathway, to promote degeneration of the distal axon fragment. Together, our results indicate that axon regeneration and degeneration are both regulated by TIR-1, and that TIR-1 carries out these two opposite functions via two divergent, and therefore separable downstream pathways. Our findings reveal a mechanism for spatial and temporal control of MAPK signaling after axon injury, which can be manipulated to shift the injury response towards repair.

## Results

### TIR-1/SARM1 inhibits axon regeneration

We asked whether TIR-1, the *C. elegans* homolog of SARM1/dSarm, regulates the injury response and found that when GABA motor axons of *tir-1* loss of function and wild-type animals were severed at the midline, significantly more axons regenerated in the absence of *tir-1* (Figure 1 B-D). We tested two putative null alleles of *tir-1*; the *qd4* allele is a 1,078 base pair deletion spanning exons 8, 9 and 10 and the *tm3036* allele is a smaller 269 base pair deletion in exon 8 that results in an early stop codon (*26, 27*). Both the *qd4* and *tm3036* deletions disrupt the C-terminal Toll/interleukin-1 Receptor (TIR) domain, which is essential for TIR-1 function (*28*) (Figure 1A). Therefore, TIR-1 inhibits regeneration of injured motor axons.

**Figure 1.**
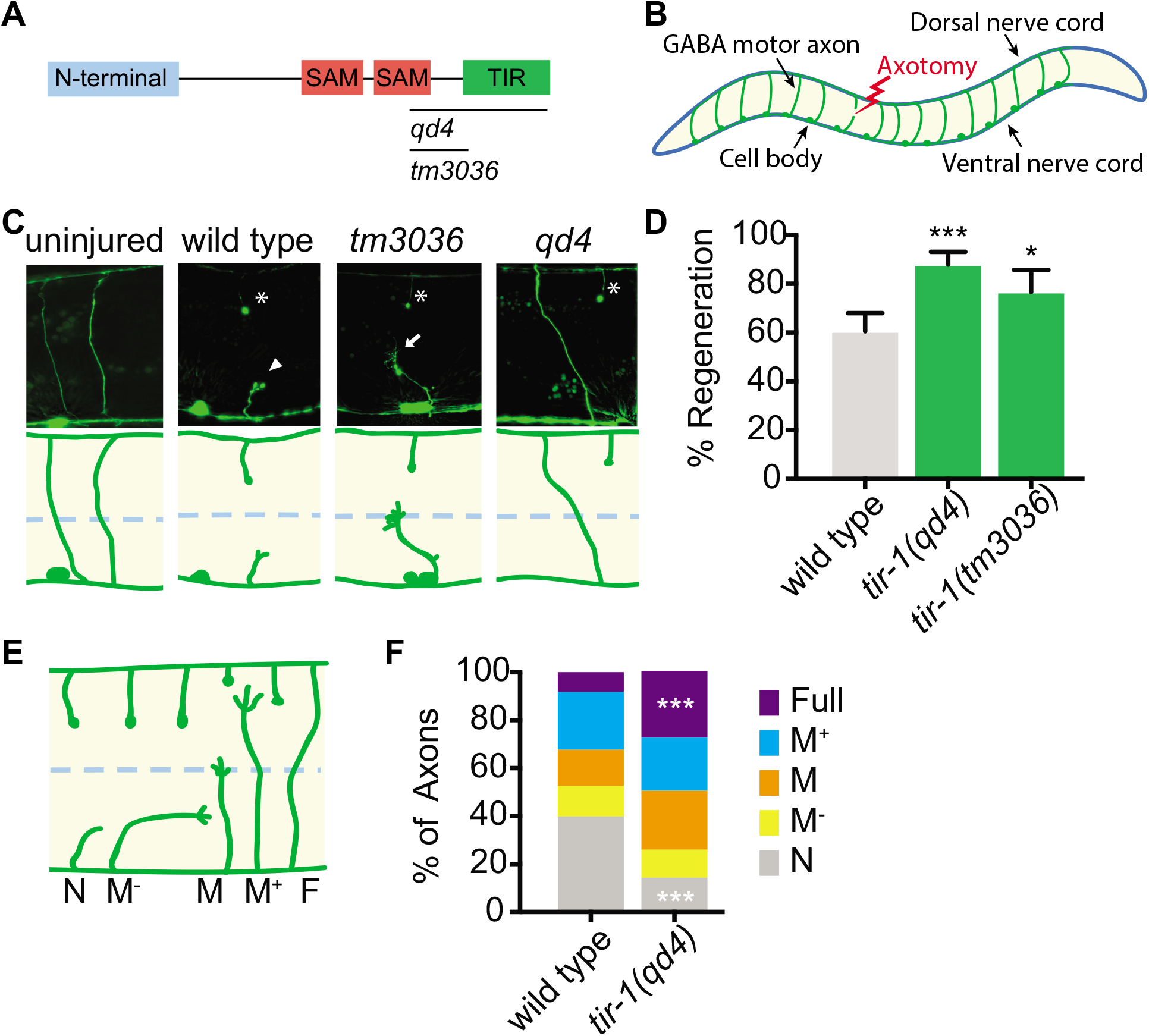
*tir-1* inhibits axon regeneration. (A) TIR-1 contains an N-terminal auto-inhibitory domain, two sterile alpha motif (SAM) domains and a toll-interleukin-1 receptor (TIR) domain. Null alleles *qd4* and *tm3036* disrupt the essential TIR domain. (B) *C. elegans* GABA motor neurons are axotomized to study the injury response in vivo and with single axon resolution. (C) Representative GABA motor neurons before and 24 hours after single laser injury at the lateral midline of the animal (dashed line). Asterisks indicate distal stumps, arrowheads indicate retraction bulbs, and arrows indicate growth cones. (D) More axons regenerate in *tir-1* mutants compared to wild-type animals, as measured by the percentage of axons that form a growth cone 24 hours after laser surgery. (E) Regeneration phenotypes were categorized according to whether regenerating axons reached landmarks M^-^, M, M^+^, F, or failed to form a growth cone, N. (F) More *tir-1*(-) axons initiate regeneration and reach the dorsal cord compared to wild-type axons, as indicated by a significant reduction in the number of axons that failed to form a growth cone (N) and a significant increase in the number of fully regenerated axons (Full). Significance relative to wild type is indicated by *p<0.05, ***p<0.001, Fisher’s exact test. Error bars represent 95% confidence intervals.

Successful regeneration of *C. elegans* motor axons includes growth cone formation, axon extension, and responsiveness to cues that guide the axon to the dorsal nerve cord. To determine which stage of the regenerative process is inhibited by TIR-1, we grouped individual injured axons into categories depending on whether it initiated regeneration and how far it regenerated towards the dorsal nerve cord (Figure 1 E, F). We found a dramatic increase in both the number of axons that initiated regeneration and in the number of axons that regenerated the full distance to the dorsal nerve cord in *tir-1* mutants compared to wild-type animals (Figure 1F). These results indicate TIR-1 inhibits the initiation of axon regeneration and that the regenerating axons are able to successfully extend and reach the dorsal cord.

*C. elegans* have one other TIR domain containing protein, which is TOL-1(*29*). To determine whether regulation of axon regeneration is a general function of TIR domain containing proteins, we compared regeneration in the presence and absence of *tol-1.* We found no significant difference between the number of axons that regenerated in *tol-1(nr2033)* null mutants and wild-type animals (Supplementary Figure 1). Therefore, TIR-1 is a TIR-domain containing protein with a specific role in axon regeneration.

### Double injury induces axon degeneration of *C. elegans* GABA motor neurons

The finding that TIR-1 inhibits axon regeneration is unexpected since SARM1 and dSarm promote the opposite response to injury, axon degeneration (*12, 13*). To determine if TIR-1 also regulates injury-induced degeneration in *C. elegans*, we needed to examine the consequences of mutating *tir-1* in manipulations that induce degeneration. Although the above axotomy assay, in which an axon is severed once, is extremely useful for investigating the genetic and cellular mechanisms that regulate axon regeneration (*30*), it is not as robust a model of injury-induced axon degeneration. The obstacle is that while proximal segments, which remain attached to the cell body, regenerate in the 24 hours following injury (*31*), the distal segments, which are no longer attached to the cell body, remain largely intact (*32*). This prompted us to develop an assay with which both injury-induced motor axon degeneration and axon regeneration can be investigated simultaneously in vivo. Here, rather than severing axons once, we severed individual axons in two places, approximately 25μM apart, which creates a middle fragment that degenerates (Figure 2A). The degeneration has similar morphological and temporal features as Wallerian degeneration, where after a brief latent period, the middle axon segment beads, fragments, and is eventually cleared (Figure 2B). We tested how reliably double injury induces degeneration of the middle fragment by quantifying the percentage of severed axons whose middle fragment either degenerated (complete clearance), partially degenerated (>65% of middle fragment degenerated) or largely remained intact (>80% of middle fragment remained). In wild-type axons, 90.2% of middle fragments degenerated 24 hours after double axotomy (Figure 2C). There was no significant difference between the number of middle fragments when visualized with either cytosolic or myristoylated and membrane bound GFP (Figure 2D). Together, this data suggests that the observed degeneration is an active process, not caused by a leaky cytosolic reporter and that the double injury reliably induces axon degeneration that can be investigated in vivo and with single-neuron precision.

**Figure 2.**
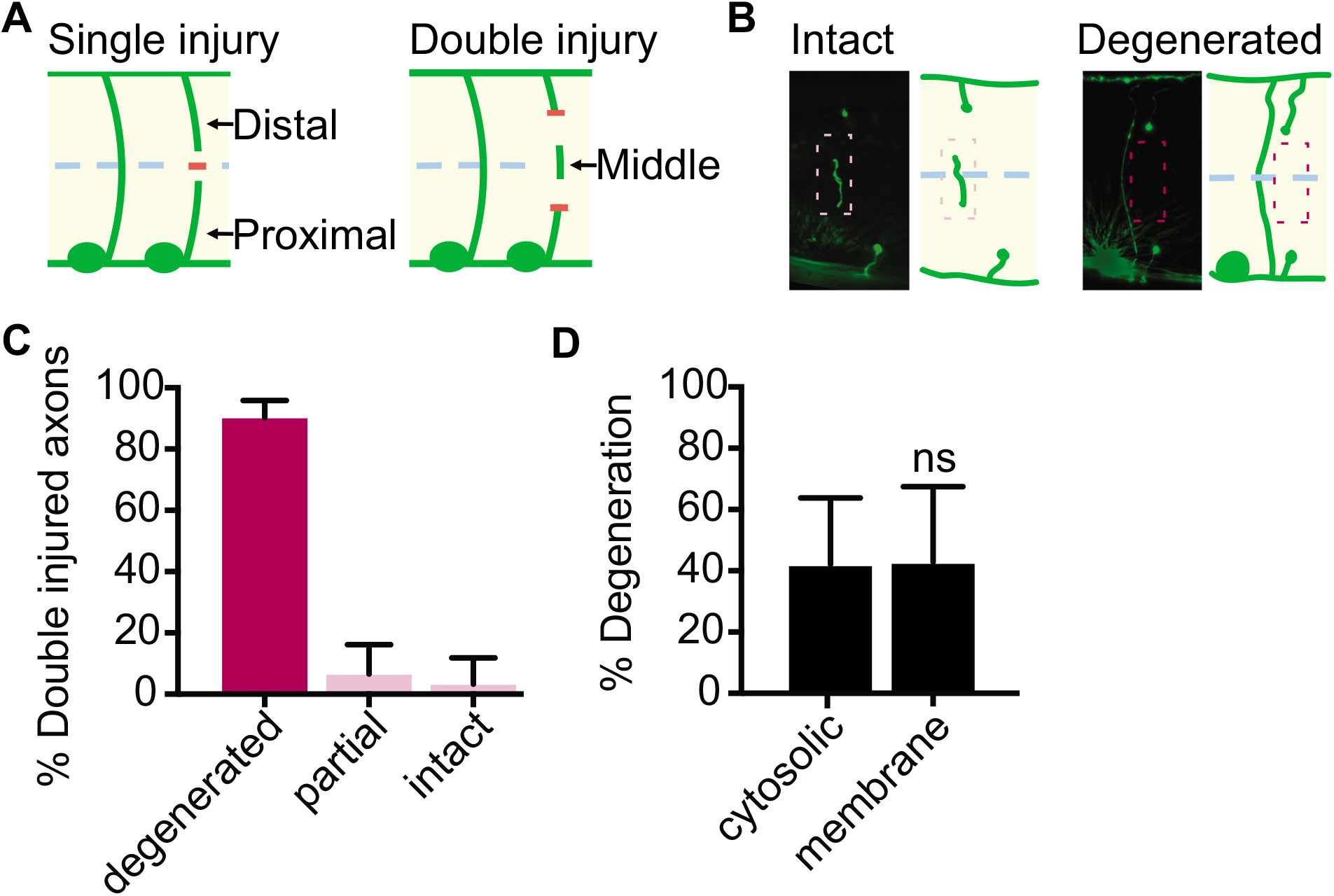
Double injury induces axon degeneration in GABA motor neurons. (A) Single and double injury models. After being severed once, axons do not degenerate. When severed twice, the middle segment degenerates. (B) Micrographs of intact and degenerated middle fragments 24 hours post axotomy. Boxes with dashed lines highlight intact (pink) and degenerated (gray) middle fragments. (C) Most middle fragments degenerate 24 hours after double injury. (D) No significant difference in middle fragment degeneration was observed between axons expressing membrane tagged- or cytosolic GFP 1.5 hours after injury, which represents an active time of degeneration at which approximately half of the severed axons are no longer visible. ns= no significance. Error bars represent 95% confidence intervals.

### TIR-1 promotes injury-induced and constitutive axon degeneration

In the absence of *tir-1*, 68% of middle fragments degenerated compared to 86% in wild-type animals (Figure 3A). In contrast, axons overexpressing *tir-1* were beaded and severely fragmented, even when not injured (Figure 3B,C,D). Upon closer examination, we found significantly less GFP expression along the axon commissures and dorsal nerve cord of *tir-1* overexpressing animals compared to non-transgenic controls (Figure 3E,F). Together, these data indicate TIR-1 promotes, but is not absolutely required for injury-induced axon degeneration. In addition, elevation of *tir-1* expression is sufficient to induce severe and chronic motor axon degeneration, even in the absence of axotomy.

**Figure 3.**
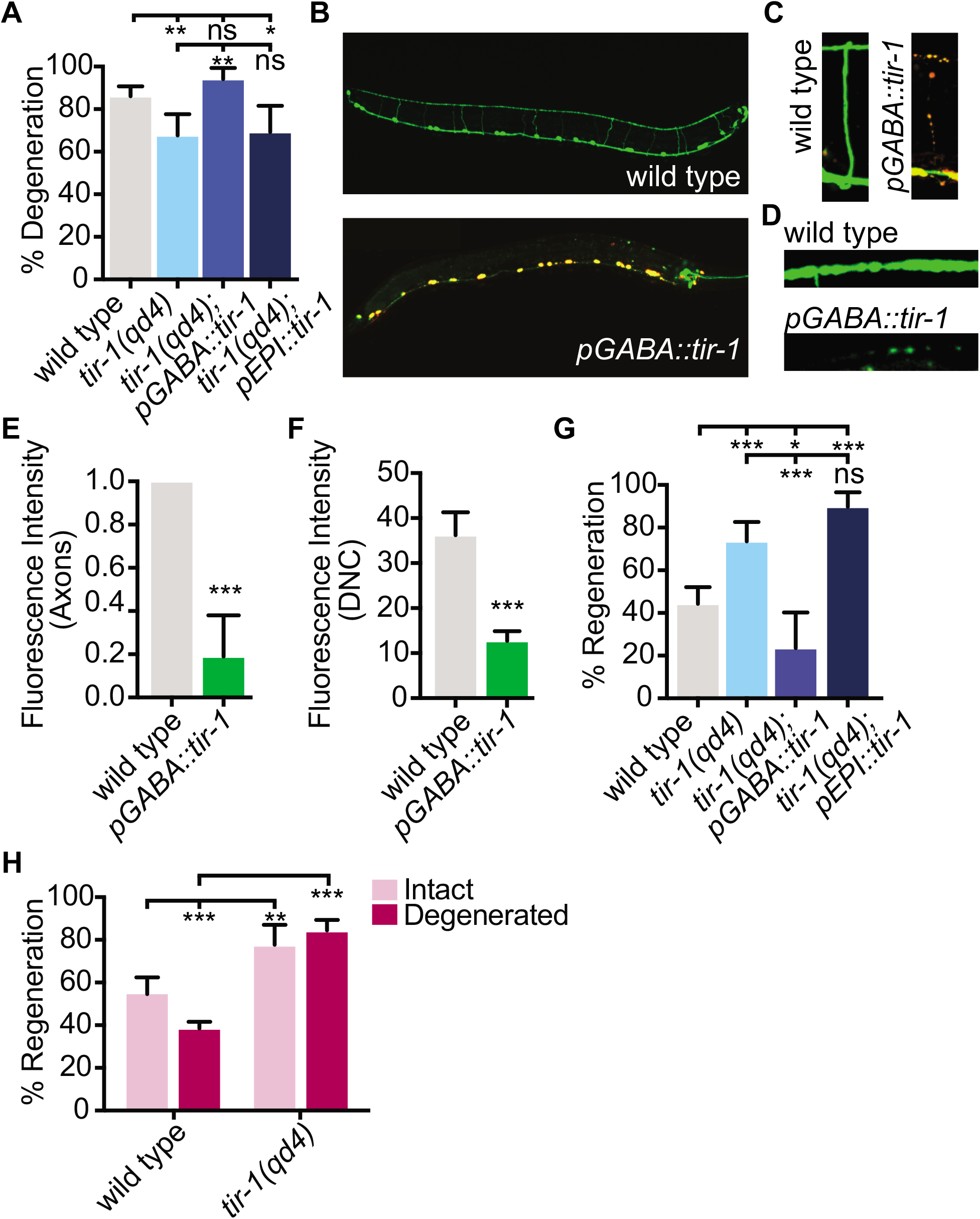
*tir-1* functions cell-autonomously to regulate the injury response. (A) GABA specific expression of *tir-1b* rescues axon degeneration in *tir-1(qd4)* mutants. (B) GABA axons seen in whole worm micrographs of wild-type animals (above) appear degenerated in transgenic animals expressing *unc-47p::tir-1::mCherry.* (C) Individual axons and (D) dorsal nerve cords of wild-type and transgenic animals expressing *unc-47p::tir-1::mCherry.* (E) Normalized GFP intensity along axon commissures and (F) GFP intensity along the dorsal nerve cord is decreased in animals that overexpress *tir-1* in GABA neurons, indicating *tir-1* promotes neuronal degeneration. (F) GABA specific expression of *tir-1b* rescues axon regeneration in *tir-1(qd4)* mutants. (G) Regeneration is inhibited when the middle fragment degenerates in wild-type animals, but not in *tir-1(qd4)* mutants. Significance relative to wild type or *tir-1(qd4)* is indicated by *p<0.05, **p<0.01, ***p<0.001. Error bars represent 95% confidence intervals.

### TIR-1 inhibits axon regeneration independently from its role in axon degeneration

Loss of *tir-1* function enhanced regeneration of axons that had been severed twice compared to wild-type axons (Figure 3G). Therefore, TIR-1 inhibits axon regeneration after double injury, as it does after single injury. To determine whether the presence of the middle fragment influenced axon regeneration we grouped axons into one of two categories 24 hours after double injury: (1) completely degenerated middle fragment or (2) intact or only partially degenerated middle fragment. We found significantly fewer wild-type axons regenerated when the middle fragment degenerated compared to wild-type axons that did not (Figure 3H). Together, these data indicate degeneration of the middle fragment inhibits axon regeneration. Of note, the number of axons that regenerated in *tir-1* mutants was similar between the injury models and regardless of whether the middle fragment had degenerated when injured twice (Figure 3H). Therefore, TIR-1 inhibits regeneration and degeneration of injured motor axons; however, its role in regeneration is not secondary to an inhibitory relationship between the two.

### TIR-1 functions cell autonomously in GABA motor neurons to regulate both axon regeneration and axon degeneration

We next asked how TIR-1 regulates axon regeneration and degeneration, and how it achieves specificity of function. TIR-1 is expressed and executes several functions in various *C. elegans* tissues including the nervous system, intestine, and epidermis (*28, 29, 33*). To determine whether TIR-1 is expressed in GABA axons, we first CRISPR tagged *tir-1* with mCherry (Supplementary Figure 2A). With this single copy integration, we observed low level fluorescence throughout the intestine, epidermis, and sheath cells of the animal, but not in GABA motor neurons. (Supplementary Figure 2B). However, in a transgenic strain expressing multiple copies of *tir-1::mCherry* under its endogenous promoter, we also observed TIR-1::mCherry expression in the cytoplasm of GABA motor neurons (Supplementary Figure 2C). Together these data indicate TIR-1 is expressed widely in tissues throughout the animal including in GABA motor neurons, where it is expressed at relatively low levels.

To determine whether TIR-1 achieves specificity of function by acting in different tissues to inhibit regeneration and promote degeneration, we asked whether TIR-1 functions in GABA axons to exclusively regulate one process or the other. We expressed *tir-1* cDNA specifically in GABA motor neurons or in the epidermis and found that GABA neuron specific expression rescued the enhanced regeneration observed in *tir-1(qd4)* mutants, while epidermis specific expression did not (Figure 3F). Similarly, GABA motor neuron specific expression of *tir-1* cDNA rescued axon degeneration in *tir-1(qd4)* mutants, while epidermis specific expression of *tir-1* did not (Figure 3A). Together, these data indicate TIR-1 functions cell autonomously in GABA neurons to both inhibit axon regeneration and promote axon degeneration after injury.

### TIR-1 regulates axon regeneration with the NSY-1 – PMK-1 mitogen-activated protein kinase signaling cascade

We next asked whether TIR-1 might direct multiple responses to injury by activating different downstream signaling events in each axon fragment. TIR-1 and SARM1/dSarm execute several cellular functions, including cleavage of NAD^+^, activation of MAPK signaling and the Axundead protein, as well as regulation of calcium signaling, gene transcription and autophagy (*14, 15, 24, 25, 28*). This informed a candidate approach to dissect the genetic pathways in which TIR-1 regulates both axon regeneration and degeneration. We found that loss of the MAP kinase kinase kinase *nsy-1/Ask1* or its downstream MAP kinase *pmk-1/p38* enhanced axon regeneration after double injury (Figure 4A,B). These regeneration phenotypes were not significantly enhanced by deletion of *tir-1*. Therefore, *tir-1* functions in the same genetic pathway as *nsy-1* and *pmk-1* to regulate axon regeneration after double injury (Figure 4A).

**Figure 4.**
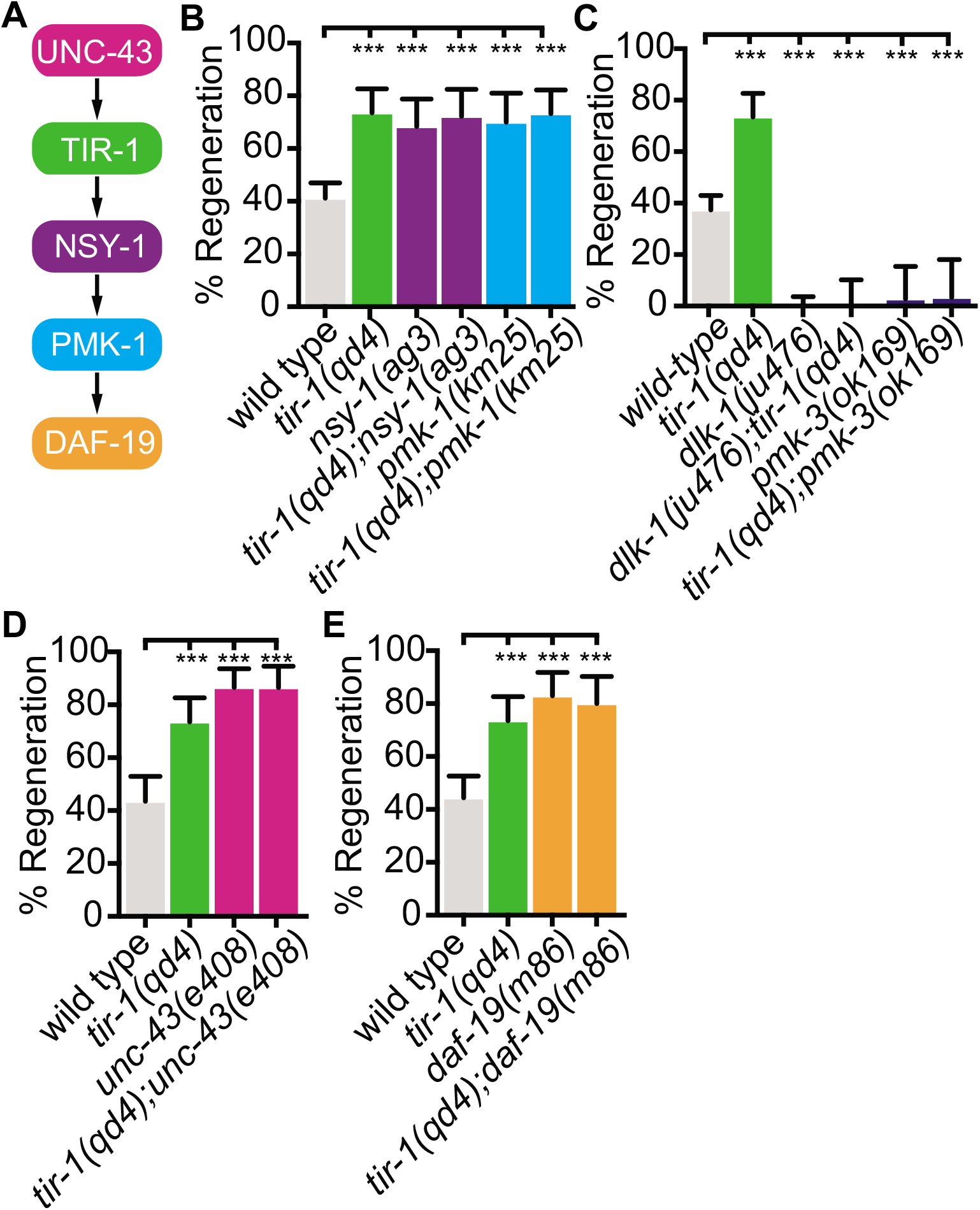
*tir-1* functions with the *nsy-1* signaling pathway to inhibit axon regeneration. (A) Components of the TIR-1— NSY-1 signaling cascade. (B) Axon regeneration is significantly increased in predicted null alleles of *nsy-1/ASK1* and *pmk-1/p38* following double injury, in both the presence and absence of *tir-1* function. Double mutants are not statistically different from single mutants. Null mutation of *unc-43/CAMIIK* enhances axon regeneration after double injury in the presence and absence of the *tir-1(qd4)* mutation. (D) Loss of the transcription factor *daf-19/RFX1-3* increases axon regeneration after double injury in the presence and absence of *tir-1.* Significance relative to wild type is indicated by *p<0.05, **p<0.01, ***p<0.001. Error bars represent 95% confidence intervals.

We also investigated whether TIR-1 regulates axon regeneration in coordination with the Dual leucine zipper MAP kinase kinase kinase DLK-1, which is essential for axon regeneration (*34, 35*). Loss of *tir-1* function did not suppress the axon regeneration phenotype of either the *dlk-1* null mutant or the downstream *pmk-3* p38 MAPK null mutant, indicating *tir-1* does not function downstream of *dlk-1* signaling (Figure 4C). As with any gene that is essential for a given function, epistasis analysis with *dlk-1* is complicated by the fact that it does not determine whether endogenous *dlk-1* function is downstream or independent from another gene. However, in combination with the finding that loss of *nsy-1* and *pmk-1* result in the same amount of regeneration as loss of *tir-1,* our data indicate that the *nsy-1–pmk-1* MAPK pathway does not act synthetically or in parallel to the *dlk-1–pmk-3* MAPK pathway to mediate *tir-1* function in axon regeneration.

Together these results indicate that TIR-1 functions with the NSY-1 – PMK-1 MAPK pathway to regulate axon regeneration.

### TIR-1 inhibits axon regeneration by regulating the RFX-type transcription factor DAF-19

During development, TIR-1 specifies the mature neuronal identity of AWC neurons downstream of the calcium/calmodulin dependent protein kinase II delta (CaMKII) ortholog, UNC-43. It does so by forming a postsynaptic protein complex with NSY-1 and UNC-43 (*28, 36*). We found that a null mutation in *unc-43(e408)* phenocopied the increased axon regeneration of *tir-1(qd4)* mutants and addition of the *unc-43(e408)* mutation to *tir-1(qd4)* mutants did not further enhance regeneration. Therefore *unc-43* and *tir-1* function in the same genetic pathway to inhibit axon regeneration (Figure 4D).

CaMKII has previously been found to regulate axon regeneration and developmental axon outgrowth, in part by regulating transcription of effector genes (*37–39*). However, the signaling pathways and set of transcription factors that mediate the effects of CaMKII activation on target gene expression during regeneration are not known. Given the genetic interactions between *tir-1*, *unc-43, nsy-1, and pmk-1* we hypothesized that UNC-43 might regulate axon regeneration via TIR-1 and NSY-1 dependent transcription. We examined this possibility by testing the regenerative ability of animals with predicted null mutations in the cAMP-dependent Transcription Factor *atf-7/ATF2/CREB5* and in the Regulatory Factor X transcription factor *daf-19/RFX,* two transcription factors regulated by *nsy-1* signaling (*24, 27, 40*). Loss of *atf-7* function did not affect GABA axon regeneration, consistent with previous studies of PLM mechanosensory axon regeneration (Supplementary Figure 3) (*41*). However, loss of *daf-19* function increased regeneration to the same extent as loss of *tir-1* and did not enhance the regeneration phenotype of the *tir-1* null mutants (Figure 4D). Together, these data are consistent with a model in which an *unc-43/tir-1/nsy-1/pmk-1/daf-19* signaling cascade regulates axon regeneration.

### TIR-1 promotes axon degeneration via a separate MAPK signaling cascade

To determine whether *tir-1* also regulates axon degeneration with the *nsy-1–pmk-1* MAPK pathway, we measured the presence or absence of degeneration 24 hours after double injury in the absence of the *pmk-1* MAPK (Figure 5A). Loss of *pmk-1* did not have a degeneration phenotype on its own and double *tir-1*; *pmk-1* null mutants phenocopy the *tir-1* single null mutant. This lack of genetic interaction indicates that *tir-1* functions independently from *pmk-1* to promote axon degeneration after injury.

**Figure 5.**
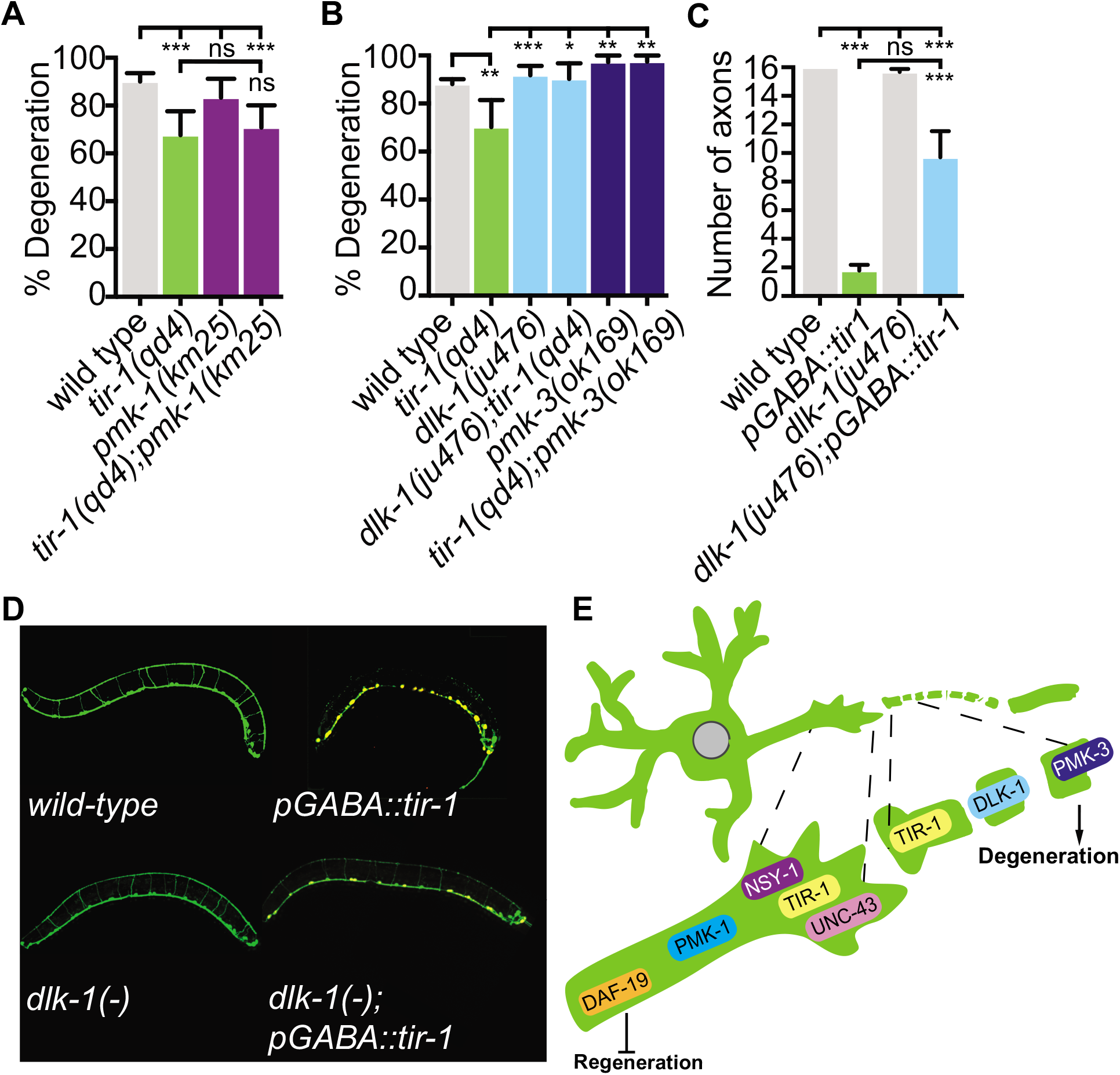
*tir-1* functions with the *dlk-1* signaling pathway to regulate axon degeneration. (A) Loss of *pmk-1/p38* does not affect degeneration with or without *tir-1* function. (B) *dlk-1/DLK/LZK* MAP3K and the downstream *pmk-3* p38 MAPK are required for the degeneration phenotype of *tir-1* mutants. (C, D) Loss of *dlk-1* suppresses the increased degeneration caused by overexpressing *tir-1* in uninjured GABA neurons. (E) TIR-1 inhibits axon regeneration and axon degeneration in the same injured axon. In the proximal stump, TIR-1 functions with the NSY-1 signaling cascade to modify gene transcription and inhibit axon regeneration. In the severed fragment, TIR-1 functions with the DLK-1 signaling cascade to promote axon degeneration. Significance relative to wild type or *tir-1(qd4)* is indicated by *p<0.05, **p<0.01, ***p<0.001. Error bars represent 95% confidence intervals.

To determine which genes do function with *tir-1* to promote axon degeneration, we returned to our candidate approach. DLK MAPK-signaling functions with SARM1/dSarm to regulate axon degeneration in mice and flies (*16, 17, 42–45*). In our assay, loss of either *dlk-1* or its downstream MAP kinase *pmk-3* did not affect axon degeneration on their own, however they did rescue axon degeneration in *tir-1* mutants (Figure 5B). This result suggests that the decreased axon degeneration phenotype of *tir-1* mutants depends on DLK-1 pathway activity. However, due to the high amount of axon degeneration in wild-type animals it is possible that there is an upper threshold in our assay that prevents us from seeing enhanced degeneration above wild-type levels. Therefore, to determine whether *dlk-1* does function with *tir-1* to regulate axon degeneration, we asked whether loss of *dlk-1* suppresses the spontaneous axon degeneration phenotype of animals that overexpress *tir-1* (Figure 5C,D). We observed an average of 16 GABA axons in wild-type animals and only 2 axons in wild-type animals that overexpressed TIR-1 (Figure 5C,D). Although *dlk-1* mutants have a wild-type number of axons, loss of *dlk-1* rescued the number of axons in the *tir-1* overexpression background (Figure 5C,D). These data indicate *dlk-1* is required for *tir-1* dependent axon degeneration. Therefore, TIR-1 regulates axon regeneration and axon degeneration with distinct MAPK signaling cascades.

## Discussion

Understanding how axons regenerate and degenerate is critical to our ability to stimulate repair after neuronal injury (*46*). Here, we identify TIR-1/SARM1/dSarm, a key regulator of axon degeneration, as a previously uncharacterized intrinsic inhibitor of axon regeneration that can be targeted to increase repair of injured motor axons. Our data indicate that to achieve this specificity of function, *tir-1* inhibits axon regeneration with the NSY-1 – PMK-1 MAPK pathway on the proximal side of the injury while it promotes degeneration with the DLK-1 – PMK-3 MAPK pathway on the distal side. Together, our findings reveal opposing mechanisms by which TIR-1 regulates the injury response.

### TIR-1/SARM1 inhibits axon regeneration independently from its role in axon degeneration

We found that TIR-1 inhibits axon regeneration and does so independently of its role in degeneration by investigating the injury response in a new model of *C. elegans* motor axon injury. *C. elegans* has been a powerful axon injury model owing to its genetic tractability, the ability to carry out experiments in vivo with single axon resolution, and its extensive genome conservation with mammals (*30, 31, 47*). In addition, the *C. elegans* motor nervous system lacks myelin and glia, enabling investigation of mechanisms that regulate axon regeneration independently of these extrinsic cues. However, *C. elegans* has not been a robust model of motor axon degeneration because for unknown reasons, when motor axons are cut once, the distal segment remains largely intact in adult animals (*32*). Instead, we found that severing an axon twice creates a middle fragment that completely degenerates and a proximal segment that is capable of regenerating. Hence, the double injury assay combines the tractability of *C. elegans* with the ability to investigate motor axon regeneration and degeneration on either side of the injury. Using this double injury assay, we found that *tir-1* both inhibits axon regeneration and promotes degeneration of injured *C. elegans* motor axons.

Either reducing the function of intrinsic inhibitors or changing the amount of extrinsic cues in the environment, such as degeneration and inflammation, can increase an injured axon’s ability to regenerate (*7–11*). Our findings indicate the enhanced regeneration in *tir-1* mutants is not a secondary consequence of reduced degeneration. First, axons lacking *tir-1* regenerate more often than wild-type axons after a single injury, even though a single injury does not cause degeneration in either wild-type or *tir-1* mutants. Second, using the double injury assay of motor axon injury in which both axon regeneration and degeneration occur, we found that *tir-1* mutant axons were as likely to regenerate in the presence or absence of degeneration on the other side of the injury. Thus, a decrease in degeneration does not account for the increased regeneration of *tir-1* mutants. Combined with our finding that *tir-1* functions within GABA axons to regulate axon regeneration, our data suggest that *tir-1* functions within the proximal stumps of injured GABA axons to regulate axon regeneration.

### TIR-1/SARM1 inhibits axon regeneration and degeneration with NSY-1 and DLK-1 MAPK signaling pathways

Our finding that *tir-1* regulates both axon regeneration and degeneration raises the question of how this single gene differentially regulates the two seemingly opposite processes in what was once the same axon. Our data indicate that TIR-1/SARM1 carries out these opposing functions by interacting with two divergent MAP kinase signaling pathways on either side of the injury. On the proximal side, TIR-1/SARM1 functions with UNC-43/CAMKII, the NSY-1/ASK1 MAPK signaling cascade and the transcription factor DAF-19/RFX to inhibit axon regeneration. The involvement of DAF-19 suggests that TIR-1 activation may change the transcriptional profile of the proximal stump into one that does not support repair. Unlike the TIR-1 pathway that regulates the innate immune response and serotonin biosynthesis, DAF-19 does not require the cotranscription factor ATF-7 to regulate axon regeneration (*40*). Therefore, TIR-1 likely achieves specificity of function by interacting with particular transcription factors in a temporal and tissue specific manner in response to injury.

On the distal side of the injury, our data suggest TIR-1 regulates degeneration with an alternate MAPK pathway. Loss of *pmk-1* function has no effect on axon degeneration, nor does it affect the degeneration phenotype of *tir-1* loss-of-function mutants, indicating TIR-1 functions independently from the NSY-1 – PMK-1 MAPK pathway to regulate axon degeneration. Instead, the DLK-1 – PMK-3 MAPK pathway is required for the *tir-1* dependent degeneration phenotype in both an injury and a constitutive degeneration model. Similarly, in mice and flies, SARM1 and dSarm interact with the signaling pathways of DLK-1 homologs Wallenda/DLK to regulate axon degeneration, indicating *tir-1* likely regulates axon degeneration, at least in part, through evolutionarily conserved mechanisms (*16–18, 21*). To date, how SARM1 ultimately leads to axon degeneration after injury is not completely clear. Besides MAP kinase signaling, effectors of SARM1 function include the downstream Axundead protein and diminished levels of NAD^+^ (*14, 15*). While there does not appear to be an Axundead homolog in *C. elegans,* the NADase activity of SARM1 and dSarm has recently been found to be conserved in TIR-1 suggesting TIR-1 may also regulate degeneration through its catalase activity (*48*).

The finding that TIR-1, a homolog of dSarm and SARM1, essential regulators of axon degeneration, also regulates the opposite response to injury, axon regeneration, is unexpected. In addition, TIR-1 was not thought to regulate injury-induced degeneration of *C. elegans* axons, likely because adult motor axons do not robustly degenerate using the single injury assay and because degeneration of injured mechanosensory axons is regulated by apoptotic engulfment machinery in surrounding tissues, and not by TIR-1 (*32, 49*). The identified dual role of TIR-1 in the injury response raises a number of compelling questions, including: Is TIR-1 differentially activated on either side of the injury to determine whether a fragment repairs itself or selfdestructs? Does TIR-1 regulate regeneration throughout the nervous system, or, like other intrinsic regulators of regeneration, does it function according to cell type and age (*50–52*)? Finally, can TIR-1 signaling be manipulated to specifically enhance axon regeneration in mammals?

In conclusion, we present a model where TIR-1 functions cell-autonomously in GABA motor neurons to regulate axon regeneration, independently from its role in axon degeneration. We show that to achieve specificity, TIR-1 directs these contrasting functions with separate downstream MAPK signaling pathways on either side of the injury. Understanding how TIR-1 regulates a balance between axon regeneration and degeneration may provide novel ways to push the nervous system towards regeneration in response to injury.

## Materials and Methods

### *C. elegans* strains and culture

Unless otherwise noted, all *C. elegans* strains were maintained at 20°C on NGM plates seeded with OP50 *E. coli.* Strains listed below were kindly provided by the Alkema, Francis, Hammarlund, Pukkila-Worley labs and the Caenorhabditis Genetics Center, which is funded by NIH Office of Research Infrastructure Programs (P40 OD010440). To visualize GABA motor neurons, strains were crossed into EG1285 (*oxIs12 [unc-47p::GFP, lin-15(+)]*). Mutants used in this study: *tir-1(qd4), tir-1(tm3036), ufIs175flp-13p::myristoylatedGFP), nsy-1(ag3), pmk-1(km25); unc-43(e408), daf-19(m86), dlk-1(ju476), pmk-3(ok169), tol-1(nr2033), atf-7(gk715*).

### Laser Axotomy, imaging, and quantification

Axotomy was performed as previously described (*49*), unless otherwise stated below. Larval stage 4 (L4) worms were mounted on 3% agarose pads and immobilized with microbeads (Polysciences) or 0.1 mM levamisole. Single injuries were made at the midline of the axon commissure and 1-3 axons were cut per worm. Double injuries were made with two targeted cuts, one roughly halfway between the ventral nerve cord and the midline of the worm and the second roughly halfway between the midline of the worm and the dorsal nerve cord, creating an approximately 25 μm middle fragment. The two injuries are performed within seconds of each other in the same order (proximal-distal). One axon was cut per worm in the double injury experiments. Axon regeneration was quantified 24 hours after injury unless otherwise stated by placing animals immobilized with microbeads or 0.1 mM levamisole on a 3% agarose pad. Animals were imaged with a Nikon 100X 1.4NA objective, Andor Zyla sCMOS camera and Leica EL6000 light source. Regeneration was scored as the percent of axons that regenerated fully to the dorsal cord (F), 75% of the distance to the dorsal cord (M^+^), to the midline (M), to below the midline or had 3+ filopodia extending from the proximal segment (M^-^). Axons that failed to regenerate, or had 2 or fewer small filopodia protruding from the proximal segment were scored as N and NF respectively.

### CRISPR

CRISPR gene editing was conducted as previously described (*53*). crRNA, tracrRNA, and Cas9 were made by IDT and injected with the rol-6 co-injection marker. The TIR-1::happylinker::mCherry crRNA sequence used is CTGCATCCACAACTTCCGAT. CRISPR mutants were genotyped and outcrossed before analysis.

### TIR-1 expression

To determine where TIR-1 is expressed we generated transgenic lines expressing TIR-1 under its endogenous promoter by injecting plasmid *bamEx102[tir-1p::tir-1b::mCherry::unc-54* 3’UTR] and co-injection marker *ceh-22::gfp* into ABC16 (*tir-1(qd4); oxIs12[unc-47p::GFP, lin-15(+)]*). L4 stage animals were imaged on Perkin Elmer Precisely UltraVIEW VoX confocal imaging system.

To determine where TIR-1 functions, we generated transgenic animals expressing TIR-1 specifically in the GABA motor neurons or in the epidermis. GABA specific expression was achieved by injecting plasmid *bamEx100[unc-47p::tir-1b::mCherry::unc-54* 3’UTR] with coinjection marker psH21 [*str-1p::gfp(+)*] into EG1285 (*oxIs12[unc-47p::GFP, lin-15(+)])* as previously described (*54*). Epidermis specific expression was achieved by injecting plasmid *bamEx101[dpy-7p::tir-1b::mCherry::unc-54* 3’UTR] with co-injection marker psH21[*str-1p::gfp(+)]* into ABC16 (*tir-1(qd4); oxIs12[unc-47p::GFP, lin-15(+)]*). The resulting transgenic strains were crossed into *tir-1(qd4)* and *dlk-1(ju476)* mutants. Axotomies were performed as described above.

### Degeneration and TIR-1 localization

Middle fragment degeneration after double injury was measured 24 hours after injury unless otherwise indicated and grouped into 3 categories: degenerated (complete clearance), partially degenerated (>65% of the middle fragment degenerated) or largely remained intact (>80% of middle fragment remained) 24 hours after double injury.

Chronic degeneration and protein localization in the absence of injury was assessed by immobilizing L4 stage worms with 0.1 mM levamisole on a 3% agarose pad. Images were taken on a Perkin Elmer Precisely UltraVIEW VoX spinning disc confocal imaging system using a 40X objective. Images were compressed and exported as TIFF files for processing in FIJI. Images were rotated and cropped to the same ROI. The axon or dorsal nerve cord segment to be analyzed was traced using a segmented line ROI and mean intensity was quantified.

### Statistical Analysis

Statistical analysis was performed using GraphPad QuickCalcs (www.graphpad.com/quickcals/) and Prism (GraphPad). Categorical data: data from repeated assays comparing a mutant and corresponding control strain were pooled for statistical analysis. Bars represent 95% confidence intervals and significance determined with Fisher’s exact test, where *p < 0.05, **p < 0.01, ***p < 0.001. Continuous data (fluorescence intensity of GFP along axon commissure) are presented as the mean with error bars representing the standard error of the mean. Significance was calculated with T-tests, where *p < 0.05, **p < 0.01, ***p < 0.001.

## Author contributions

V. Julian contributed to experimental design, investigation, interpretation, and manuscript preparation. A. Byrne contributed to experimental design, interpretation, and manuscript preparation. We thank A. Zeamer for technical contribution to Figure S2, W. Joyce and M. Liu for construction of reagents, along with M. Alkema and V. Budnik for helpful comments on the manuscript.

## Competing interests

Authors have no competing interests.

## Data and materials availability

All data is available in the main text or the supplementary materials. All materials used in the analysis are available upon request.

## Supplementary Figures

**Supplementary Figure 1.**
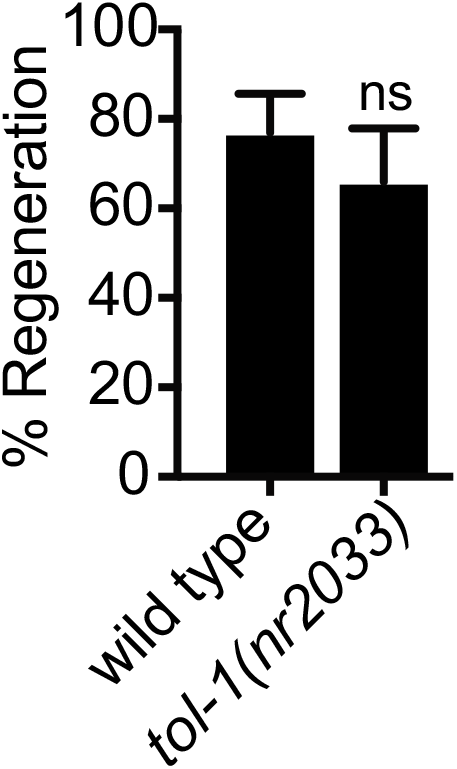
tol-1 does not regulate axon regeneration. (A) Axon regeneration is not significantly different in *tol-1(nr2033)* mutants compared to wild-type controls 24 hours after laser surgery. Error bars represent 95% confidence intervals.

**Supplementary Figure 2.**
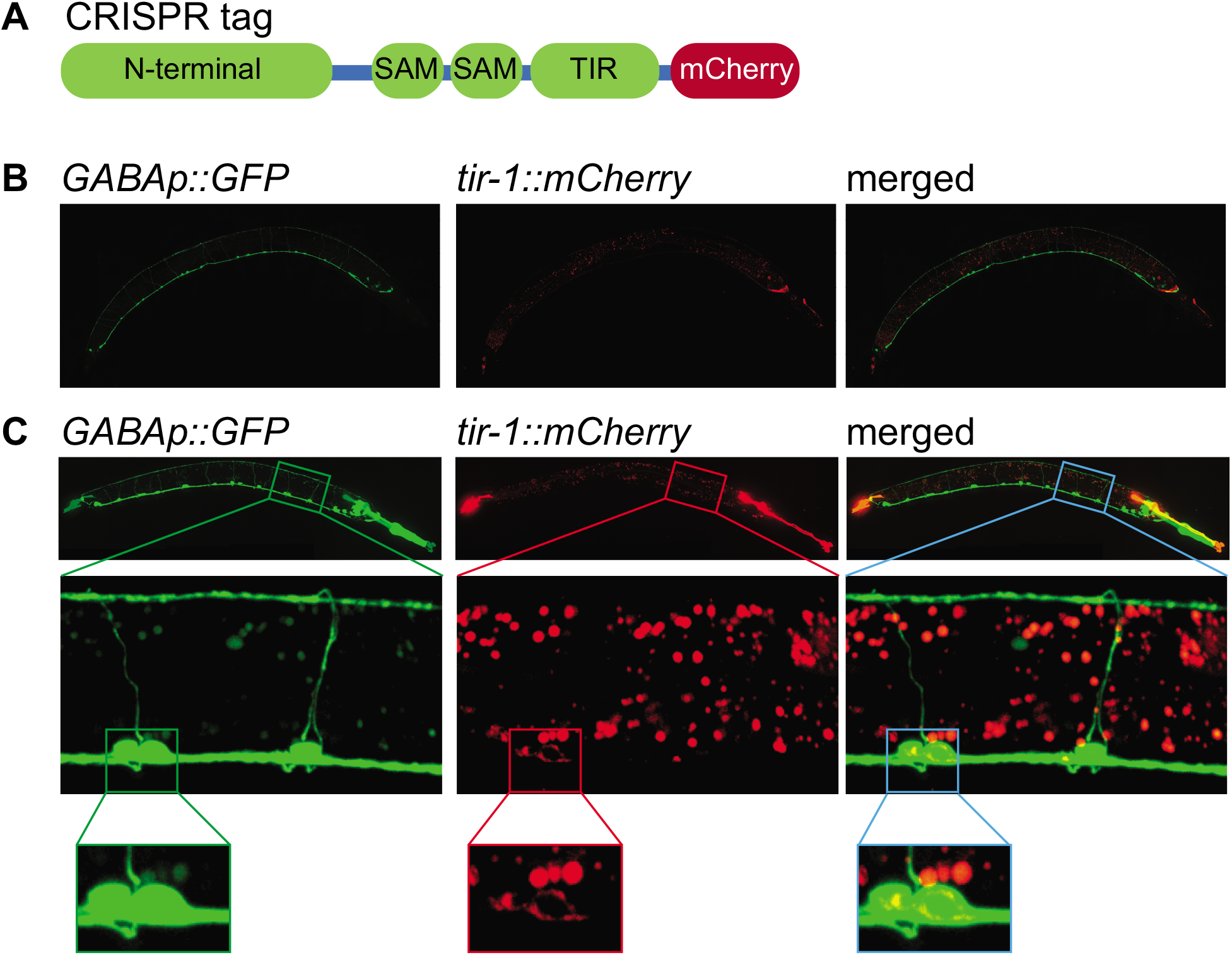
TIR-1 localizes to GABA motor neurons. (A) The *tir-1* gene was CRISPR tagged with mCherry. (B) Representative whole worm micrographs of animals expressing endogenous *tir-1::mCherry.* No overlap was seen with the GABA specific GFP marker *unc-47p::GFP.* (C) Representative micrographs of animals expressing a *tir-1b::mCherry* extrachromosomal array and the GABA specific GFP marker *unc-47p::GFP.* TIR-1b::mCherry was observed in cell bodies of GABA neurons (enlarged boxes).

**Supplementary Figure 3.**
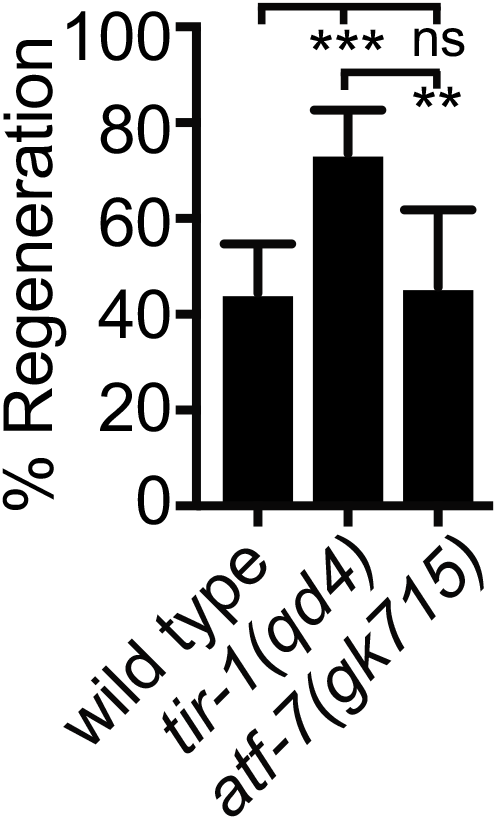
The *atf-7* transcription factor does not regulate axon regeneration. (A) Axon regeneration is not significantly different between *atf-7(gk715)* mutants and wild-type controls 24 hours after laser surgery. Significance relative to wild type or *tir-1(qd4)* is indicated by **p<0.01, ***p<0.001. Error bars represent 95% confidence intervals.

## Notes

### Competing Interest Statement

The authors have declared no competing interest.

